# Nutri Complex 150+: a new and effective approach to facial rejuvenation

**DOI:** 10.1101/2024.10.05.616805

**Authors:** Anna Privitera, Greta Ferruggia, Martina Contino, Salvatore Maugeri, Massimo Zimbone, Venera Cardile, Giuseppe Caruso, Maria Violetta Brundo

**Affiliations:** Department of Drug and Health Sciences, University of Catania, Catania, Italy; Department of Biomedical and Biotechnological Sciences, University of Catania, Italy; Department of Biological, Geological and Environmental Science, University of Catania, Catania, Italy; National Research Council of Italy - Institute for Microelectronics and Microsystems (CNR-IMM), Catania, Italy; Unit of Neuropharmacology and Translational Neurosciences, Oasi Research Institute-IRCCS, Troina, Italy

**Keywords:** AMPLEX plus, Aging, Facial rejuvenation, Growth factors, Cytokines

## Abstract

Skin is the largest multifunctional human organ and possesses a complex multilayered structure with the ability to regenerate and renew. The key role in skin regeneration is played by fibroblasts, also playing an important role in wound healing process. We used different methods to evaluate on human fibroblasts the *in vitro* effects of a new compound called Nutri Complex 150+ (NC150+), containing a mixture of 20 different biologically active factors (GF20) and exosomes isolated and purified from bovine colostrum. NC150+ was able to significantly enhance cell proliferation/metabolic status of fibroblasts at both 24 and 48 hours compared to untreated (control) cells. NC150+ was also able to enhance the ability of human fibroblasts to close the wound scratch. Our findings demonstrate the ability of NC150+, based on a new technology called AMPLEX plus, to enhance cell proliferation/metabolic status of fibroblasts. The obtained results also suggest how NC150+ could be potentially effective in treating skin injury.

## Introduction

Aged skin is characterized by appearance of wrinkles, laxity, and pigmentary irregularities. These changes occur under the influence of extrinsic (e.g., UV exposure, smoking, air pollution) and intrinsic factors, like cellular changes [1,2]. Between the two, skin aging due to intrinsic factors is inevitable and strongly depend on time progresses, and changes in intrinsically (chronologically) aged skin can produce morbidity in the aging population [3].

Skin is an important component of outward beauty and is the focus of various cutaneous surgical and non-surgical procedures [4]. As people age, the structure and function of the epidermis and dermis are altered. The epidermal permeability barrier is largely determined by the quality and quantity of growth factors such as epidermal growth factor (EGF) and keratinocyte growth factor (KGF) secreted by keratinocytes, the most abundant cell type in the epidermis. In the aged epidermis, the cells divide more slowly, and the level of skin growth factors reduced parallel to a decline in keratinocytes proliferation and an increase in apoptosis of these cells leading to thinning epidermis [5,6].

During the aging process, the dermis also undergoes significant alterations. Unlike the epidermis, which is made up of dense keratinocytes, the dermis is comprised primarily of extracellular matrix (ECM) components, including collagen, which is the major component of ECM, and elastic fibers. Aging negatively impacts primary fibroblasts proliferation, which is significantly reduced [7]. As a result, collagen production is decreased, and collagen fibers become fragmented and coarsely distributed. Other ECM components, including elastin fibers, glycosaminoglycans (GAGs), and proteoglycans (PGs), also change during aging, ultimately leading to a net functional dermal component’s deficiency [8]. As this network loosens and unravels with time, aging features emerges on skin surface, contributing to increase skin fragility [9].

Growth factors and cytokines, directly influences the biosynthesis of collagen, elastin and ECM components, but their binding and signaling decreases with age. It should be noted that there is no single molecule involved in skin rejuvenation [10]. Therefore, successful repair of damaged skin and collagen synthesis require the balanced involvement of growth factors and cytokines [11]. These biologically active factors are secreted in the form of nanosized extracellular vesicles (EVs) by cells. Based on size and release mechanism, different types of vesicles can be distinguished: exosomes are the smaller particles which are released not directly from the cell membrane, but via the fusion of the multivesicular bodies (MVBs) and the plasma membrane [12,13].

Growth factors and cytokines carried by exosomes alter target cell bioactivities, such as increased cell proliferation, cell migration and protein release [14]. Thus, providing these factors from outside through specific products with regenerative action can help reduce the appearance of fine lines, wrinkles and improve skin tone and texture.

Skin changes that result from the chronologic aging process prompt patients to seek procedures to correct all changes and achieve rejuvenation of the skin. Changes in the facial skeleton impact significantly on facial appearance and their correction has constituted one of the main focuses of facial rejuvenation procedures [15,16]. Studies have already investigated the potential beneficial effects of colostrum-derived exosomes on reducing skin damage, promoting the proliferation of epidermal cells and, additionally, the production of collagen type I by fibroblasts [17]. Our work proposes a new compound Nutri Complex 150+ (NC150+) that used a new technology called AMPLEX plus, capable of stimulating skin regeneration and revitalization. It is a strategy resulting from the combined action of a mixture containing 20 different biologically active factors, especially rich in growth factors, essential for specific function (GF20), and concentrated exosomes, previously purified and isolated from bovine colostrum. Exploiting the combined action of GF20 and colostrum derived exosomes, could improve the regenerative and repairing effect on skin deficiencies.

## Materials and Methods

Materials and reagents used to carry out the experiments related to the present work, were all of analytical grade and supplied by Thermo Fisher Scientific Inc. (Pittsburgh, PA, USA) or by Sigma-Aldrich Corporate (St. Louis, MO, USA), if not otherwise specified.

### Preparation of GF20

GF20 is a colostrum-derived solution composed, as described by [19], of growth factors and cytokines, which are involved in tissue regeneration. GF20 was obtained following the procedure proposed by [20]. Briefly, the colostrum was initially diluted 1:10 in deionized water with NaCl. Subsequently, the suspension was centrifuged at 12,400 x g at 20 to 25 °C to remove the fat fraction. The supernatant was ultrafiltered to remove large proteins like casein, and in the last step of dialysis, the obtained product was subjected to sterilizing cross-filtration. Immunoglobulin removal was performed by the tangential filtration or cartridge filters.

### Isolation and characterization of colostrum-derived exosomes

Isolation of exosomes from colostrum was performed by serial centrifugations to remove all debris and molecules not relevant for our study. Initially, colostrum was centrifuged at 2000 x g for 10 min. Subsequently, only the intermediate part was collected, while the superficial part consisting of fat and the pellet (debris) were discarded. The collected sample was ultracentrifuged at 10,000 x g for 30 min. The pellet with the debris was discarded, while the supernatant was ultracentrifuged at 100,000 x g for 70 min to isolate the exosomes. At the end of ultracentrifugation, the pellet containing the exosomes was resuspended in PBS and centrifuged again at 100,000 x g for 70 min to wash and purify the exosomes. Characterization, performed by light scattering measurements, showed that exosomes derived from colostrum have a spherical shape with a diameter of 150 ± 1.2 nm and a concentration of 4.2 × 10^12^ particles/mL [19].

### Nutri Complex 150+ composition

Nutri Complex 150+ contain 5 mL of AMPLEX plus that includes 20 billion exosomes and 200 mg of GF20. The exosomes in AMPLEX plus are loaded passively with growth factors and cytokines transferred from GF20.

### Propagation and maintenance of cells

Human non immortalized fibroblasts were cultured in Dulbecco’s Modified Eagle Medium (DMEM) supplemented with 10% FBS, streptomycin and penicillin (0.3 mg mL^−1^ and 50 IU mL^−1^, respectively), and GlutaMAX (1 mM) by using 25 or 75 cm^2^ polystyrene culture flasks. Cells were maintained in a humidified environment (37 °C, 95% air/5% CO_2_), and split every 2–3 days depending on cell confluence.

### Analysis of cell proliferation/metabolic status

One day before the experiment, fibroblasts were harvested by using a trypsin-EDTA solution, counted with a C-Chip disposable hemocytometer, and plated in 96-well plates (4 or 8 × 10^3^ fibroblasts/well). The following day, fibroblasts were treated with NC150+ (1%) and incubated for 24 and 48 hours in a humidified environment (37 °C, 5% CO_2_). At the end of the treatment, cell proliferation/metabolic activity were measured by the well-known MTT assay [21,22]. Briefly, at the end of the treatment, MTT solution (1 mg/mL in DMEM medium) was added to each well and cells were incubated for 2 hours in a humidified environment (37 °C, 5% CO_2_). During the final step, DMSO was used to melt the crystals, while the Synergy H1 Hybrid Multi- Mode Microplate Reader (Biotek, Shoreline, WA, USA) was used to read the absorbance at 569 nm. Values were normalized with respect to control untreated fibroblasts and were expressed as the percent variation of cell proliferation/metabolic activity.

### Scratch-wound assay

Human non immortalized fibroblasts were plated into six-well plates at the density of 2.5 × 10^6^ cells/well allowed to incubate until confluence. Once the confluence was reached, cells were scraped by using a 1000 μL sterile pipette tip in order to form a wound [23]. Cells were washed by using PBS to remove detached cells before adding the medium, in absence or presence of NC150+, used at different percentages (up to 1%). Among the different percentages tested, 0.1% was the one selected, being the best in terms of activity and viability during the entire period of 96 hours. Untreated cells were considered as control. At 0 (T0), 24 (T24), 48 (T48), 72 (T72) and 96 (T96) hours’ time points, images of the wound area in the same field were captured by using Nikon ECLIPSE Ts2 – FL inverted microscope (Nikon Instruments Inc., Melville, NY, USA; magnification x4).

### Statistical analysis

The statistical analysis was carried out by using Graphpad Prism software (version 8.0) (Graphpad software, San Diego, CA, USA). Two-way analysis of variance (ANOVA), followed by Tukey’s post hoc test, was used for multiple comparisons. The statistical significance was set at *p*-values < 0.05. Data were reported as the mean ± SD of at least 2 independent experiments.

## Results

### Characterization of colostrum exosomes

The dynamic light scattering (DLS) revealed the size of particle, while the morphology was verified by SEM (Figure 1). The average diameter size was 10^7^ ± 1.2 nm and exosomes presented a spherical morphology. The concentration mean was 3.0 × 10^12^ particles/mL.

**Figure 1.**
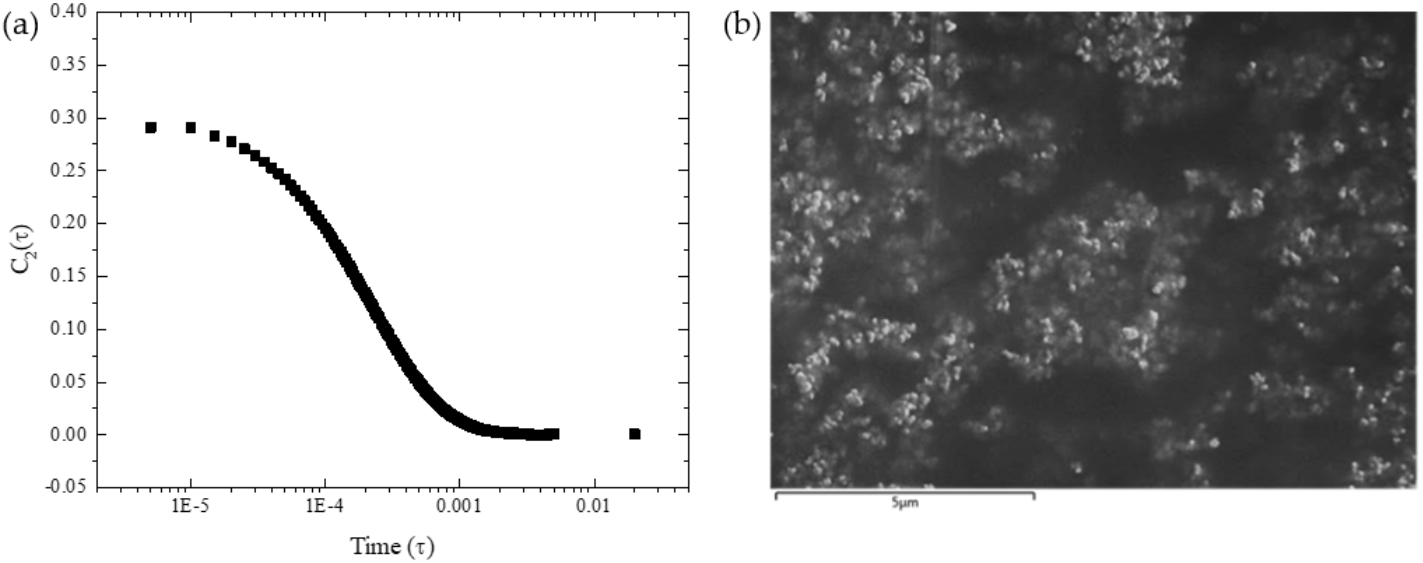
Characterization exosomes from colostrum. (a) Analysis of size by DLS; (b) Observation of exosomes morphology by SEM.

### Nutri Complex 150+ possess the ability to induce cell proliferation in human fibroblasts

**Figure 2**. depicts the effects of NC150+, a compound that used the AMPLEX plus technology, on cell proliferation/metabolic status of fibroblasts. NC150+ was able to significantly increase (p < 0.001) cell proliferation/metabolic status of fibroblasts at both 24 and 48 hours compared to untreated cells (control) (Figure 2). Interestingly, 24 hours treatment with NC150+ produced an inductive effect higher than that observed for the same treatment at 48 hours (Figure 2).

**Figure 2.**
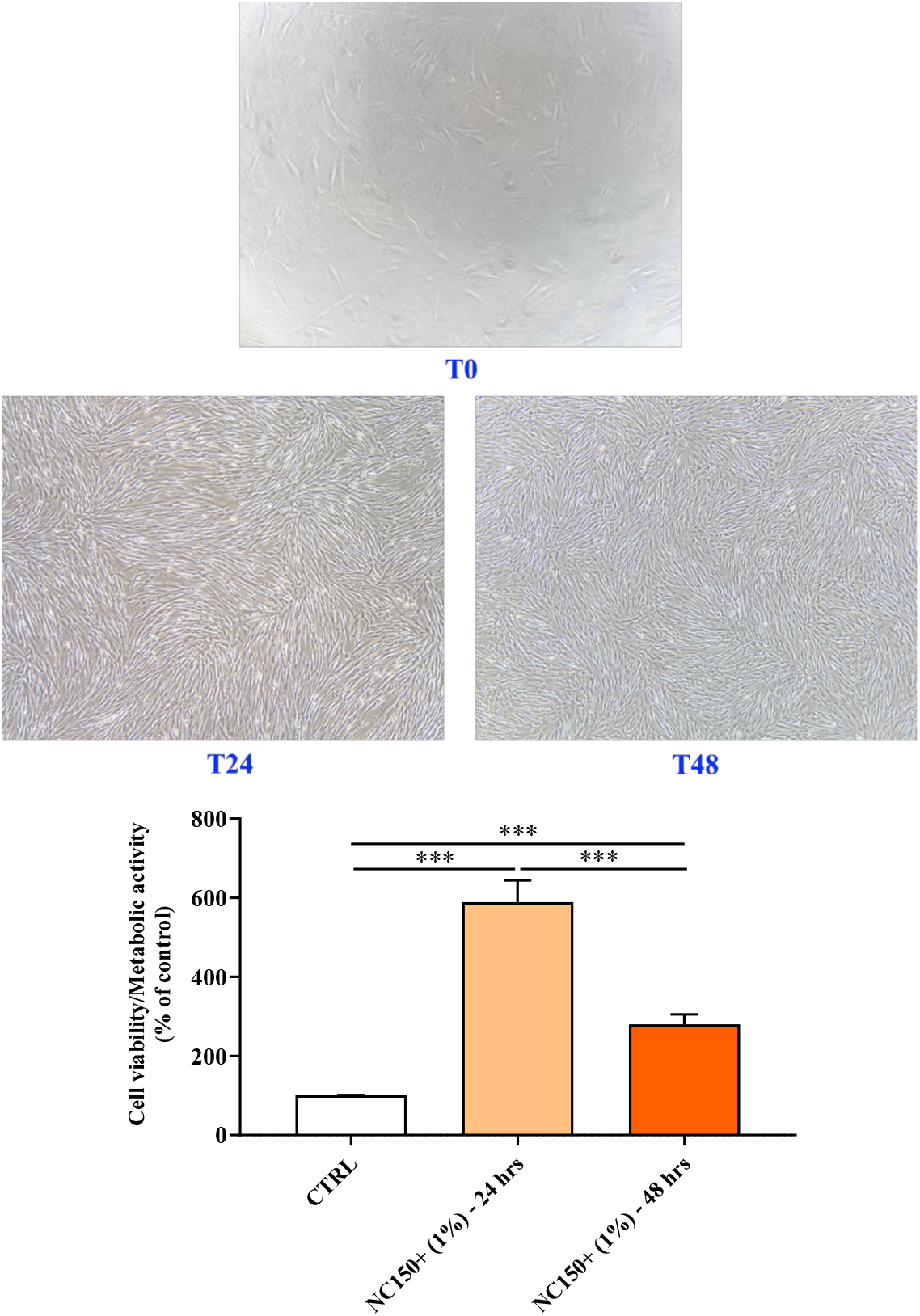
Cell proliferation/metabolic activity in untreated and treated with Nutri Complex 150+ (1%) (NC150+) for 24 and 48 hours assessed by MTT assay (8 × 10^3^ fibroblasts/well). Data were the mean of six values and were expressed as the percent variation with respect to the cell proliferation/metabolic activity recorded in untreated (CTRL) cells. Standard deviations were represented by vertical bars. Significantly different, \*\*\**p* < 0.001.

### Nutri Complex 150+ enhances the ability of human fibroblasts to close the wound scratch

Figure 3 reported the effects of NC150+ on wound scratch closure carried out by human fibroblasts during a total of 96 hours. As expected, based on the results observed by employing the MTT assay, the treatment of human fibroblasts with NC150+ for 24 hours (T24) increased the number of migrating cells detectable in the wound site compared to untreated cells. The difference between the two experimental conditions was still remarkable at T48, with the wound scratch almost completely closed in cells treated with NC150+. After 72 hours, cells treated with NC150+ reached complete confluence, while 96 hours were needed in the case of untreated fibroblasts.

**Figure 3.**
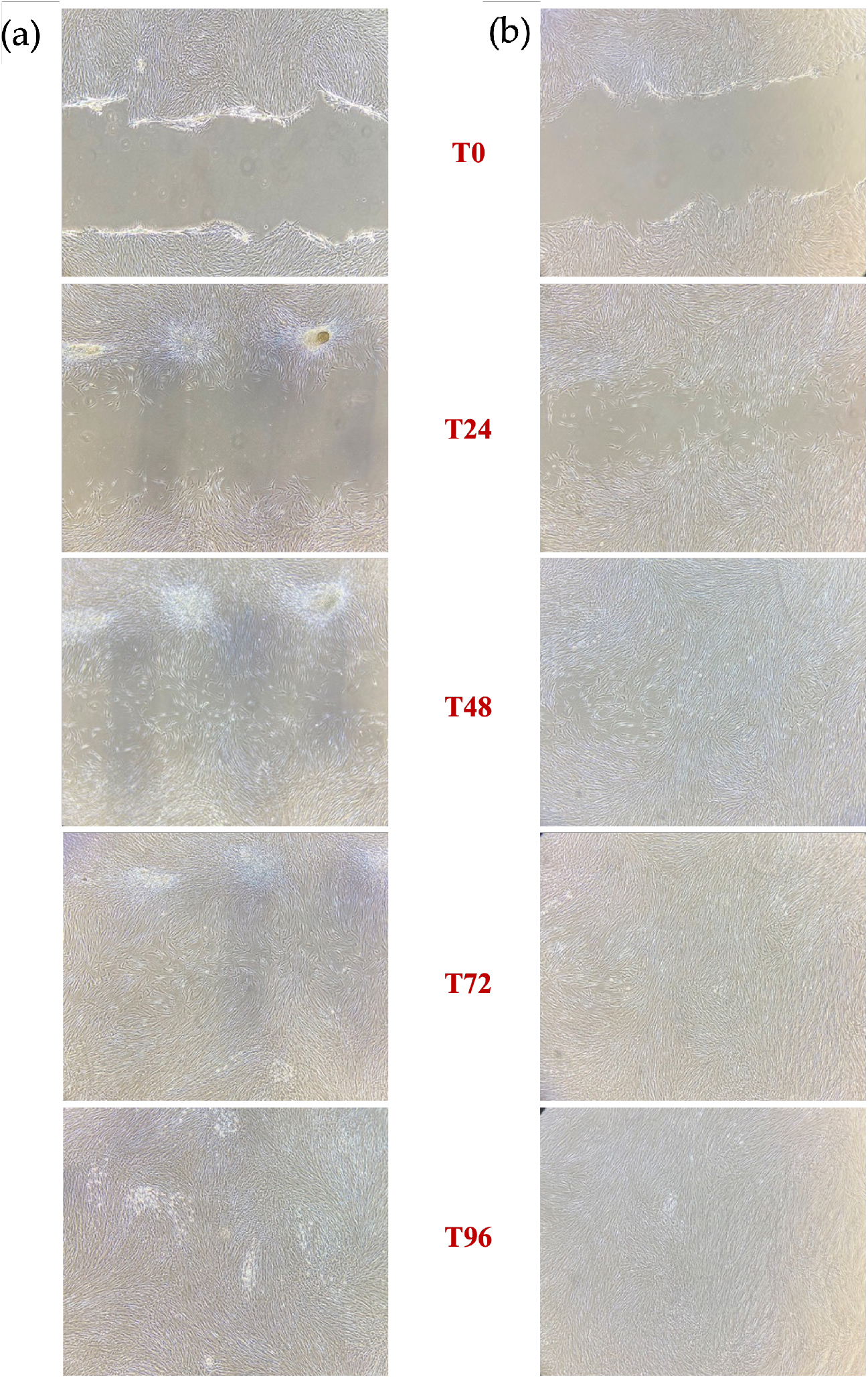
Representative images of wound scratch closure carried out by (a) untreated human fibroblasts and (b) human fibroblasts treated with NC150+ (0.1%) at 0 (T0), 24 (T24), 48 (T48), 72 (T72), and 96 (T96) hours’ time points. Untreated cells were considered as a control.

## Discussion

Currently, most research on skin changes with age focuses on the undesirable aesthetic aspects of skin aging, but the deterioration of the skin with age is more than just an aesthetic problem that affects most people, both male and female [24]. Growth factors and cytokines are responsible for maintaining skin homeostasis, which helps to keep skin alive and healthy. They are protein molecules that significantly support the repair of damaged skin, as a result of aging, and several studies, over the past 15 years, have highlighted the potential benefits of growth factors, showing improvements in the appearance of wrinkles, texture, fine lines and discoloration [25-29]. In particular, growth factors and cytokines present in GF20 may be involved in many processes. TGF-β, whose concentration in GF20 is 3.5 ± 0.15 pg/mL [19], controls an immense number of cellular responses and figures prominently in development and homeostasis of must human tissues, such as bone and cartilage. TGF-β acts as a chemoattractant for monocytes and is described as the most potent chemoattractant for neutrophils [30, 31]. It also increases epithelial cell migration at the edges of wounds and stimulates the collagen secretion by fibroblasts [32]. IGF-1 has multiple important biological activities. It belongs to a class of multiple peptide growth factors involved in cell proliferation, differentiation, and individual growth, especially in cartilage and muscle tissue [31]. IGF-1, whose concentration in GF20 is 3.1 ± 0.08 pg/mL [19], is also an anabolic growth factor that is known to improve the metabolic rate and to promote protein synthesis. The metabolic effects of IGF-1 are correlated to GH hormone, in fact IGF-1 functions as the major mediator of growth hormone (GH) [33,34]. The bFGF – FGF2 is one of the strongest vascular growth factors involved in the process of proliferation, differentiation, and survival of a wide variety of cell types [35]. It is described as a multipotent growth factor that regulates cell growth as well matrix composition, chemotaxis, adhesion, and migration of epithelial cells [36]. bFGF concentration in GF20 is 3.9 ± 0.49 pg/mL [19]. VEGF is known to play an essential role in vascular development. It stimulates endothelial cell (EC) proliferation and migration and promotes the growth of new blood vessels during embryonic development, new blood vessels after injury and new collateral vessels to bypass obstructed vessels. VEGF stimulates wound healing through angiogenesis, but likely promotes collagen deposition and re-epithelialization as well [37,38]. VEGF concentration in GF20 is 3.3 ± 0.42 pg/mL [19]. The EGF, whose concentration in GF20 is 2.4 ± 0.36 pg/mL [19], is a common mitogen factor that contributes to proliferation, differentiation, and growth of different types of cells, especially epithelial cells, and fibroblast. EGF is not only capable of potent mitogenic effects, but also stimulates angiogenesis, influencing wound closure and epidermal repair [39,40]. PDGF regulates many cellular processes, including cell proliferation, migration, and angiogenesis. It stimulates chemotaxis of fibroblasts and smooth muscle cells and promotes collagen synthesis by fibroblast. These properties suggest that PDGF plays an important role in wound repair process, significantly helping the healing wound [41]. PDGF concentration in GF20 is 2.7 ± 0.35 pg/mL [19]. The KGF is a member of fibroblast growth factor family (FGF-7) that shows numerous activities. KGF is a potent mitogen for different types of epithelial cells, which regulates migration and differentiation of these cells, especially keratinocytes. KGF is present in epithelialization – phase of wound healing during which helps the up-regulation of VEGF and increases transcription of factors involved in the detoxification of ROS preserving keratinocytes [42,43]. KGF concentration in GF20 is 2.6 ± 0.40 pg/mL [19]. HGF induces epithelial cell proliferation, motility, and morphogenesis in various tissues. These pleiotropic function of HGF is essential for the construction of normal tissue architecture [44]. HGF is also known angiogenesis factor by its property to promote endothelial cell growth, survival, and migration [45]. HGF concentration in GF20 is 3.4 ± 0.15 pg/mL [19]. GM-CSF, whose concentration corresponds to 3.9 ± 0.38 pg/mL [19], increases re-epithelialization, recruiting neutrophils, monocytes and lymphocytes and it stimulates phagocytosis in matured leucocytes. In addition, GM-CSF increases angiogenesis and lymphangiogenesis [46,47]. G-CSF influences the proliferation, differentiation, and survival of haemopoietic stem cells and mature neutrophils and induces the proliferation of endothelial cells and their migration to the damaged area postinjury, accelerating would healing. G-CSF also stimulates angiogenesis and neurogenesis [48,49]. The concentration in GF20 is 2.8 ± 0.27 pg/mL [19]. CCL11 is a chemokine that stimulates eosinophil chemotaxis and enhances their adhesion to endothelial cells, contributing to the activation of eosinophils recruited into the damage tissue and playing roles in epithelial remodeling. It modulates fibroblast activities by increasing their proliferation and collagen synthesis and promotes angiogenesis [50,51]. Its concentration in GF20 corresponds to 2.4 ± 0.37 pg/mL [19]. TNF−α regulates the activity of fibroblasts, vascular endothelial cells and keratinocytes and promotes synthesis of extracellular matrix proteins, which are closely involved in the healing of injured tissues. TNF-α induces angiogenesis and shows anti-malignant cell cytotoxicity, especially in combination with Interferon. It contributes to bone remodeling by activating osteoclasts and plays an important role in controlling infection [52,53]. In GF20, the concentration is 2.3 ± 0.35 pg/mL [19]. NGF promotes fibroblast and keratinocyte proliferation, extracellular matrix component expression and secretion and it stimulates the proliferation and differentiation of local immune cells, blood vessels and even neurite outgrowth. NGF contributes also to the reestablishment of a normal sensory and sympathetic innervation of the damaged skin [54,55]. The concentration is equal to 2.6 ± 0.83 pg/mL [19]. INF-γ promotes macrophage activation and mediates antiviral and antibacterial immunity. During cutaneous wound healing, INF-γ contributes to angiogenesis and collagen deposition and encourages CD4+ T helper polarization, which contributes to the initial wound microenvironment. INF-γ controls cellular proliferation and apoptosis and it also coordinates lymphocyte-endothelium interaction [56,57]. In GF20, its concentration is 3.3 ± 0.38 pg/mL [19]. BMP-2 induces the formation of both cartilage and bone. BMP-2 directs the development of neural crests cells into neural phenotypes and induces chemotaxis, mesenchymal cell proliferation and differentiation. BMP-2 enhances angiogenesis and regulates the healing process by promoting dermal and epidermal growth, leading to keratinized and thickened skin [58,59]. Its concentration corresponds to 3.1 ± 0.27 pg/mL [19]. SDF1-α is one of the most potent chemokines for stem cells recruitment. Its concentration corresponds to 3.3 ± 0.30 pg/mL [19]. SDF1-α can regulate multiple physiological processes such as organogenesis, regeneration and angiogenesis and administered at the site of ischemia, is associated with ischemic neo-vascularization. SDF1-α has an important role during bone fracture healing and play a central role during the process of angiogenesis, such as chemotaxis, cell proliferation, migration, and the secretion of angiopoietin factors [60,61]. IL-2 stimulates proliferation and enhances function of T-cells, NK cells and B-cells and induces B- cells to generate secretory IgM. IL-2 stimulates macrophages to gain maturity and elaborate TGF-β. In synergy with bFGF, IL-2 can impact the rate and quality of the closure at the wound site by promoting skin cell proliferation. Furthermore, it can promote angiogenesis [62,63]. Its concentration in GF20 is equal to 3.3 ± 0.30 pg/mL [19]. IL-4 has an important role in regulating antibody production, hematopoiesis, inflammation, and T cells development and can rescue B-cells from apoptosis, enhancing their survival and promoting their secretion of IgE and IgG. It is almost twice as potent as TGF- β at stimulating collagen synthesis by fibroblasts and is a potent mitogen for microvascular endothelial cells [64,65]. Its concentration is 0.9 ± 0.05 pg/mL [19]. IL-6 stimulates the activation of osteoclasts for the continual physiological process of bone remodeling as well as for the repair process during bone healing. IL-6 induces excess production of VEGF, leading to enhanced angiogenesis and increased vascular permeability and can aid keratinocyte proliferation and the generation of collagen by dermal fibroblasts [66,67]. Its concentration is equal to 1.8 ± 0.30 pg/mL [19]. IL-17A promotes the production of G-CSF which act in synergy with TNF-α to induce neutrophil recruitment. It can stimulate macrophages and neutrophils to produce lactoferrins and regenerating proteins, helping to kill bacteria. IL-17A maintains integrity of the mucosal tissues and promotes wound closure, myofibroblast differentiation and collagen deposition [68]. In GF20, its concentration is 3.0 ± 0.47 pg/mL [19].

In recent year, also exosomes with their high stability and direct stimulation of target cells, have demonstrated potential as anti-aging treatment for skin defects [69]. Exosomes express characteristics of the cell from which they are released and contain several molecules (such as peptide, growth factors), functioning as paracrine molecules that interact with the ECM and adjacent cells [70] or distant cells [71]. To exploit their capabilities, effective loading strategies for exosomes have been developed. Exosomes are loaded through two processes: exogenous and endogenous. In the exogenous or direct loading process, molecules are loaded onto the purified exosomes after isolation from cells. This is further subdivided into active and passive loading. Passive loading, as happens in AMPLEX plus technology, involves loading the cargo into exosomes through diffusion, while active loading is characterized by the disruption of exosome membranes [72].

The aim of this study was to overview the revitalizing effects on the skin of NC150+, obtained by the combination of GF20 bioactive factors and colostrum-derived exosomes. GF20. NC150+ is composed of a high variety of growth factors, cytokines and other protein molecules known to act synergistically. Among the characterized factors, bFGF and TGF-β showed the highest concentrations. These growth factors are crucial in providing a substantial stimulation of fibroblasts, contributing to facial rejuvenation.

Aging is accompanied by a decrease in skin function, mainly regenerative, and, therefore, a decrease in the skin’s ability to renew and maintain proper homeostasis. This involution of the skin is primarily dependent on the changes associated with the activity of fibroblasts, representing the main cell population of the skin [73]. The activity of these cells is also fundamental for the regulation of numerous physiological processes such as brain development and wound healing [74].

We have demonstrated that NC150+ was able to significantly enhance cell proliferation/metabolic status of fibroblasts. Since fibroblasts also play a key role during wound healing process [77-80], the last set of experiments was carried out to investigate the potential effects of NC150+ in accelerating the closure of an induced wound scratch. As expected, based on the results regarding cell proliferation/metabolic status, the treatment of human fibroblasts with NC150+ strongly increased the number of migrating cells detectable in the wound site compared to untreated cells at each time points. Of note, cells treated with NC150+ closed the wound scratch completely at 72 hours, while 96 hours were needed in the case of untreated fibroblasts.

## Conclusions

The findings reported in the present study demonstrate the ability of NC150+, based on AMPLEX plus technology, to enhance cell proliferation/metabolic status of fibroblasts. The obtained results suggest how NC150+ could represent a beneficial treatment to repair damaged skin and to restore and maintain skin health and youth.

## Author Contributions

Conceptualization, G.C. and M.V.B.; methodology, A.P., G.F., M.C., S.M., and M.Z.; software, A.P., M.Z., and G.C.; validation, A.P. and G.F.; formal analysis, A.P., G.F., M.C., S.M., and M.Z.; investigation, A.P. and G.F.; resources, V.C., G.C., and M.V.B.; data curation, A.P., M.Z., G.C., and M.V.B.; writing—original draft preparation, G.C and M.V.B.; writing—review and editing, A.P., G.F., V.C., G.C., and M.V.B.; supervision, G.C. and M.V.B.; funding acquisition, M.V.B. All authors have read and agreed to the published version of the manuscript.

## Institutional Review Board Statement

This study was performed in line with the principles of the Declaration of Helsinki and does not require approval by the Ethics Committee of University of Catania.

## Informed Consent Statement

Not applicable.

## Data Availability Statement

The data presented in this study are available on request from the corresponding author.

## Acknowledgments

The authors would like to thank the International PhD Program in Neuroscience at University of Catania (A.P.) and PhD program FSE Notice 1/2021 (G.F.) at the University of Catania (Italy). Authors also acknowledge the PON project Bio-nanotech Research and Innovation Tower (BRIT), financed by the Italian Ministry for Education, University and Research (MIUR) (Grant no. PONa3_00136). The research team likes to express its sincere gratitude to Dermoaroma Italy srl for the generous provision of the NC150+ and related consumables that facilitated the successful completion of this research project.

